# Targeting lncRNA *JINR1* with programmable Circular Active Nano DNAzyme (CANDe) suppresses Japanese Encephalitis Virus infection

**DOI:** 10.64898/2026.05.30.728920

**Authors:** Chandrika Sharma, Suryansh Sengar, Dharani Sen, Vivek Sharma, Souradyuti Ghosh

## Abstract

Oligonucleotide therapeutics such as antisense oligonucleotides (ASOs) and small interfering RNAs (siRNAs) enable sequence-specific gene silencing but rely on endogenous cellular machinery and often require extensive chemical modification for stability and efficacy. DNAzymes offer a mechanistically distinct alternative through intrinsic catalytic RNA cleavage; however, their therapeutic translation has been limited by nuclease susceptibility, structural constraints, and synthetic challenges. Here, we report the development of Circular Active Nano DNAzyme (CANDe), an enzymatically synthesized circular DNAzyme platform designed to enhance stability without backbone modification. The therapeutic potential of CANDe constructs was investigated against Japanese Encephalitis Virus (JEV) infection associated host long-noncoding RNA *JINR1* (LINC01518). CANDe constructs were generated via splint-assisted ligation and incorporate modular elements, including catalytic cores (8–17 or 10–23), target-binding arms, and structural stems. Circularization conferred marked resistance to exonuclease-mediated degradation compared to linear DNA, maintaining structural integrity under nuclease-rich conditions.,CANDe targeting the lncRNA *JINR1* achieved effective JINR1 knockdown in SHSY-5Y with and without JEV infection. This was accompanied by reduced expression JEV RNA and titers. In line with this, CANDe constructs attenuated of virus-induced cytotoxicity and apoptosis. Among the constructs, 10–23-based CANDe targeting the JINR1-1 site exhibited the strongest overall activity. These findings establish CANDe as a modular, modification-free DNAzyme platform that combines catalytic efficiency with enhanced stability, enabling effective host-directed antiviral intervention. This approach highlights topological engineering as a viable alternative to chemical modification for advancing DNAzyme-based therapeutics.

## Introduction

Oligonucleotide-based therapeutics have emerged as powerful tools for sequence-specific gene modulation, including antisense oligonucleotides (ASOs; e.g., nusinersen, inotersen), small interfering RNAs (siRNAs; e.g., patisiran, givosiran), microRNA mimics, and aptamers (e.g., pegaptanib)^1–8^. These modalities predominantly rely on endogenous cellular machinery, such as RNase H1-mediated RNA degradation for ASOs and RNA-induced silencing complex (RISC)–dependent mRNA cleavage for siRNAs and microRNAs, to elicit therapeutic effects ^9–11^. Although chemical modifications (e.g., 2′-O-methyl sugars and phosphorothioate linkages) enhance target specificity and nuclease resistance, dependence on host proteins such as Argonaute-2 or RNase H can limit efficacy under conditions of cellular stress, infection, or impaired protein function^10,12^.

In contrast, DNAzymes (deoxyribozymes) catalytic motifs such as 10–23 and 8–17, identified through in vitro selection of, autonomously catalyze RNA cleavage at purine– pyrimidine junctions via a metal-ion mechanism involving coordinated nucleophilic attack, independent of cellular machinery^13–15^. This intrinsic catalytic activity enables multiple turnover reactions, with reported catalytic efficiencies (k_cat_/K_M_) approaching 10^9^ M^−1^ min^−1^, surpassing the largely stoichiometric modes of action of ASOs and siRNAs^13^. In addition, DNAzymes can be readily redesigned by simple rearrangement of the substrate-binding arms and have demonstrated promising pre-clinical activity against oncogenic, and inflammatory targets, with stability further improved through locked nucleic acid or 2′-fluoro modifications^16^. Collectively, their machinery-independent activity and robustness across diverse physiological conditions position DNAzymes as a compelling alternative for gene silencing in cellular environments where host protein function is compromised.

Despite their catalytic prowess, DNAzyme advancement into therapeutics is stymied by inherent design constraints. The extended catalytic core (~15-30 nt) plus substrate arms totals 40-60 nt, far longer than ASOs/siRNAs (~20 nt), complicating automated phosphoramidite synthesis: yields plummet below 20% for >50-mers, escalating costs >10-fold and producing toxic byproducts like acrylonitrile from deprotection^17–19^. Secondary structures, like G-quadruplexes or stem-loops in the metal ion-binding core, sterically hinder uniform backbone modifications (PS, 2’-Ome, XNA)^20–25^. These structural constraints severely limit the incorporation of backbone or sugar modifications; even minor alterations (e.g., phosphorothioate or 2′-O-methyl substitutions) can disrupt the catalytic core geometry and reduce activity by over 90%. In contrast, antisense oligonucleotides (ASOs) tolerate extensive chemical optimization, enabling enhanced stability and prolonged half-lives extending to several days^26^. The resulting lack of modifiability limits DNAzyme stability (t_1_/_2_ ~hours) and tissue penetration, restricting them largely to local delivery^21,27,28^. Circularization offers a compelling alternative: topological locking confers exonuclease resistance (t_1/2_ >100-fold), unfolds dynamically for substrate docking, and biophysically stabilizes the construct without modifications, retaining 25-80% activity in 10-23 circles. Pioneering work demonstrated circular DNAzymes cloned via M13/T4 ligase targeting β-lactamase mRNA^29^. However, conventional ligation protocols often yield low product fractions (frequently <10–20%) due to competing intermolecular ligation, multimerization, and structural constraints, limiting scalability toward GMP production^30,31^. Splint-templated or aptamer-guided methods improve specificity but require bespoke selection, limiting versatility. These synthetic bottlenecks, compounded by core rigidity, have relegated DNAzymes to niche research, despite superiority over linear oligonucleotides in autonomy and potency. Streamlined, high-yield circularization sans modifications remain essential for clinical translation.

Despite the implementation of regional vaccination programs, JEV continues to exert a profound socioeconomic burden across South and Southeast Asia, manifesting in approximately 68,000 clinical cases annually^32–35^. The virus is characterized by a high fatality rate of 20–30%, and for survivors, the prognosis is often accompanied by permanent neurological sequelae^35–38^. Recent transcriptomic analyses have identified host lncRNAs, lncRNA *JINR1* (LINC01518), as an attractive target for mitigating JEV infection^39–41^. Mechanistic studies reveal that *JINR1* modulates the expression of GRP78, a crucial endoplasmic reticulum chaperone that facilitates viral entry and subsequent replication^42^. When *JINR1* expression is suppressed,GRP78 levels are reduced, leading to the significant attenuation of viral titers and a reduction in virus-mediated cell death. Although *JINR1* knockdown has been achieved using locked nucleic acid (LNA)–modified antisense oligonucleotides (ASOs), the high cost, synthetic complexity, and limited scalability of LNAs limit their applicability in regions where JEV is endemic, motivating the development of more accessible oligonucleotide platforms with comparable specificity.

In this study, we report the development of a *JINR1* targeted circular DNAzyme-based nanoconstruct platform, termed Circular Active Nano DNAzyme (CANDe), synthesized through a splint-assisted enzymatic circularization strategy designed to enhance intracellular stability and functional persistence. CANDe is organized into discrete structural elements, including a splint-binding region for ligation, a stabilizing stem, a catalytically active DNAzyme core (8–17 or 10–23), and flanking target-binding arms that enable sequence-specific RNA binding and cleavage. This enzymatic assembly strategy eliminates the need for synthetic backbone modifications, thereby minimizing the toxicity of phosphorothioate or analogus modifications while preserving catalytic activity. Using JEV-infected SH-SY5Y neuronal cells as a model system, we demonstrate efficient CANDe-mediated cleavage of the lncRNA *JINR1* transcript, accompanied by reduced expression of its downstream effector GRP78 and a concomitant decrease in viral RNA levels and plaque-forming units. Together, these results establish CANDe as a modular DNAzyme nanoplatform for targeted gene silencing.

## Materials and Methods

### General

All oligonucleotides used in this study (sequences listed in Table 1) were purchased from Eurofins Genomics and supplied following high-purity salt-free (HPSF) purification. The oligonucleotides were used without further purification. T4 polynucleotide kinase (M0201S), T4 DNA ligase (M0202SVIAL), Exonuclease I (M0293S), Exonuclease III (M0206SVIAL), bovine serum albumin (BSA, P8108S), adenosine 5′-triphosphate (ATP, P0756L), T4 DNA ligase buffer (B0202SVIAL), and NEB buffer 1 (B7001SVIAL) were obtained from New England Biolabs (USA). Tris base (71033), spermidine (17000), and dithiothreitol (DTT, 17315) were purchased from Sisco Research Laboratories Pvt. Ltd. (SRL), India. Phosphodiesterase I from Crotalus adamanteus venom (P3243-1VL) was procured from Sigma-Aldrich. Fetal bovine serum (FBS), Dulbecco’s Modified Eagle Medium (DMEM), DMEM/F12 medium, Oxoid phosphate-buffered saline (BR0014G), and paraformaldehyde (Q23996) were purchased from Thermo Fisher Scientific. Penicillin–streptomycin solution (9802000) was obtained from Invitrogen. Agarose MP (11388983001) and crystal violet (V5265) were acquired from Sigma-Aldrich. All aqueous solutions were prepared using ultrapure water from a Milli-Q Type II water purification system (Millipore). Experimental procedures were carried out using a Prima-96 thermal cycler (HiMedia), a SpectraMax iD5 microplate reader (Molecular Devices), and a CFX Opus 96 Real-Time PCR system (Bio Rad).

### Phosphorylation

Precursor linear ssDNA oligonucleotides were phosphorylated as per our existing protocol41. Briefly, phosphorylation was carried out in the presence of T4 PNK, ATP, phosphorylation buffer (50 mM Tris-HCl pH 7.5, 10 mM MgCl_2_, 1 mM ATP, 10 mM DTT), DTT, spermidine, and water to make up the volume. The solution was incubated at 37°C for 3 h, followed by inactivation by heating at 75°C for 20 min. Afterwards, the splint was mixed with the 5′-phosphorylated padlock, followed by annealing.

### Ligation

Circularization of the annealed 5′-phosphorylated DNA was carried out in the presence of T4 DNA ligase, ATP, and T4 DNA ligase (50 mM Tris-HCl pH 7.5, 10 mM MgCl_2_, 1 mM ATP, 10 mM DTT) buffer. The ligation mixture was then incubated at room temperature (25°C) for 4 h, followed by enzyme inactivation at 75°C for 20 min.

### Exonuclease Digestion

The exonuclease digestion was performed in a 100 µL reaction volume, as described below. For exonuclease digestion, the reaction was carried out with both the exonuclease enzymes in exonuclease III buffer (10 mM Bis-Tris-Propane-HCl, 10 mM MgCl_2_, 1 mM DTT, pH 7). All the reactions were supplemented with DTT (1 mM) to ensure complete digestion. The reagents were added while keeping the vials on ice. The solutions were incubated at 37°C for 4 h, followed by enzyme inactivation at 85°C for 20 min. The sample concentration was quantified by densitometric analysis of sample bands in 12% Urea PAGE to confirm the final product’s concentration.

### Snake venom phosphodiesterase (SVPD) assay

250 ng of linear DNA or CANDe were incubated with SVPD (final 2 mU/μL) in DMEM 10% FBS media at 37°C for time points of 0, 5, 15, 30, 60, and 90 minutes. The reaction was terminated by heating the sample at 80°C for 5 min. Sample stability was determined by densitometric analysis of sample bands, along with an untreated 250 ng control, in 12% Urea PAGE.

### Cell culture

SH-SY5Y cells were cultured in DMEM/F12 medium supplemented with 10% heat-inactivated fetal bovine serum (FBS), 2 mM L-glutamine, and penicillin–streptomycin (Gibco) under standard conditions (37°C, 5% CO_2_). PS cells were maintained in Dulbecco’s modified Eagle medium (DMEM) supplemented with 10% heat-inactivated fetal bovine serum (FBS), 2mM of glutamine, and penicillin/streptomycin (Gibco) at 37°C and 5% CO_2._.

### Virus propagation and Plaque Assay

The GP78 strain of JEV was propagated in Vero cells as described previously^41,42^. Briefly, Vero cell monolayers were inoculated with JEV at 0.1 MOI and incubated at 37°C and 5% CO_2_ for 72 hpi (hours post-infection) until the infected monolayers showed the cytopathic effect. Subsequently, the culture supernatant was harvested by centrifugation at 2,000 rpm for 20 min at 4°C, followed by filtration using syringe filters with 0.8/0.2 µm (Acrodisc Syringe Filters, #4658) and aliquoted to generate a virus stock. JEV titer was determined by plaque assay in PS cells. All virus stocks were aliquoted and stored at −80°C.

The plaque asssay was done using PS cells. For plaque assays, 2 × 10^5^ PS cells were seeded into 6-well culture plates. After 24 h, the monolayers were infected with serial dilutions of cell culture–derived JEV (GP78) for 2 h. After adsorption, PS cells were washed twice with PBS to remove unbound virus, then overlaid with a 1:1 mixture of 2% low-melting agarose and growth medium. Cultures were incubated for 4 days at 37°C in 5% CO_2_, after which they were fixed in 4% formaldehyde for 1h at 37°C. Plaques were visualized by staining with 0.5% crystal violet solution for 5 min, followed by three washes with RO water. Viral titers were determined as plaque-forming units (PFU) per millilitre according to the formula: PFU = (N × DF)/V, where *N* is the number of plaques, *DF* the dilution factor, and *V* the inoculum volume.

### CANDe Transfection

SH-SY5Y cells were reverse-transfected with CANDe using Lipofectamine RNAiMAX (Invitrogen, #13778-075) according to the manufacturer’s protocol. Briefly, 0.8×10^6^ cells were reverse-transfected with 60 pmol CANDe and followed by virus infection at 18 hours post-transfection.

### Viral infection

One-day post-seeding, SH-SY5Y cells were serum starved for 2 h and infected with JEV at an MOI of 5 in an incomplete DMEM-F12 medium at 37°C for 2 h. After virus adsorption, the inoculum was removed, and the cells were washed with PBS. After removing the virus-containing medium, the cells were grown in growth media with reduced serum (5%) for 36h, followed by RNA extraction.

### Cell Proliferation Assay

Cell proliferation was evaluated using the WST-1 reagent (Roche, #05015944001) according to the manufacturer’s instructions. Briefly, SH-SY5Y cells were seeded into 96-well plates and subjected to reverse transfection with Circular Scrambled JINR1-S (control), CANDe 8-17 JINR1-1, CANDe 10-23 JINR1-1 and CANDe 10-23 JINR1-2 at a final concentration of 60 pmol. At the indicated timepoints, the cell proliferation was measured by recording the absorbance at 450nm.

### Caspase Assay

Caspase-3/7 activity was measured using a luminometric assay kit (Promega, #G8090) according to the manufacturer’s protocol. SH-SY5Y cells were transfected with Circular Scrambled JINR1-S (control), CANDe 8-17 JINR1-1, CANDe 10-23 JINR1-1 and CANDe 10-23 JINR1-2 and subsequently luminescence was recorded at the indicated time point to measure the relative caspase 3/7 activity.

### RNA isolation and real-time PCR

Total RNA was isolated using the Direct-zol RNA Miniprep Kit (Zymo, #R2050). Subsequently, 1 µg of RNA was reverse transcribed into cDNA with the PrimeScript First-Strand cDNA Synthesis Kit (Takara, #6110A). Quantitative real-time PCR (qRT-PCR) was carried out using the SYBR Green PCR Kit (Takara, #RR820A) on a CFX Opus 96 Real-Time PCR system (Bio Rad)Each reaction was performed in triplicate, with GAPDH serving as the internal reference gene. Relative expression levels were determined using the 2^−ΔΔCt^ method.

## Results

### Design consideration of CANDe

The circular nano-construct CANDe was designed to enable efficient RNA cleavage by DNAzymes without requiring chemical modification, while providing resistance to nuclease degradation. In addition, the construct is not sequestered by proteins, allowing it to dissociate after catalysis and participate in multiple turnover events. For target selection, we employed the well-characterized RNA-cleaving DNAzymes 8–17 and 10–23 against lncRNA JINR1, chosen for their dependence on Mg^2+^, a physiologically abundant cofactor (Figure 1A). The 8– 17 DNAzyme contains a 14 nt catalytic core that cleaves RNA at AG dinucleotides, whereas the 10–23 DNAzyme comprises a 15 nt core targeting purine–pyrimidine junctions (AU, AC, GU, or GC), thereby expanding the range of accessible cleavage sites. Cleavage sites were identified within regions targeted by the validated LNA ASOs JINR1-1 and JINR1-2 to ensure structural accessibility for hybridization (Figure 1B). Binding arms were subsequently designed flanking cleavage sites located centrally within these regions, with 8 nt lengths selected to balance target binding affinity with efficient product release and catalytic turnover (Figure 1C–E).

**Figure 1.**
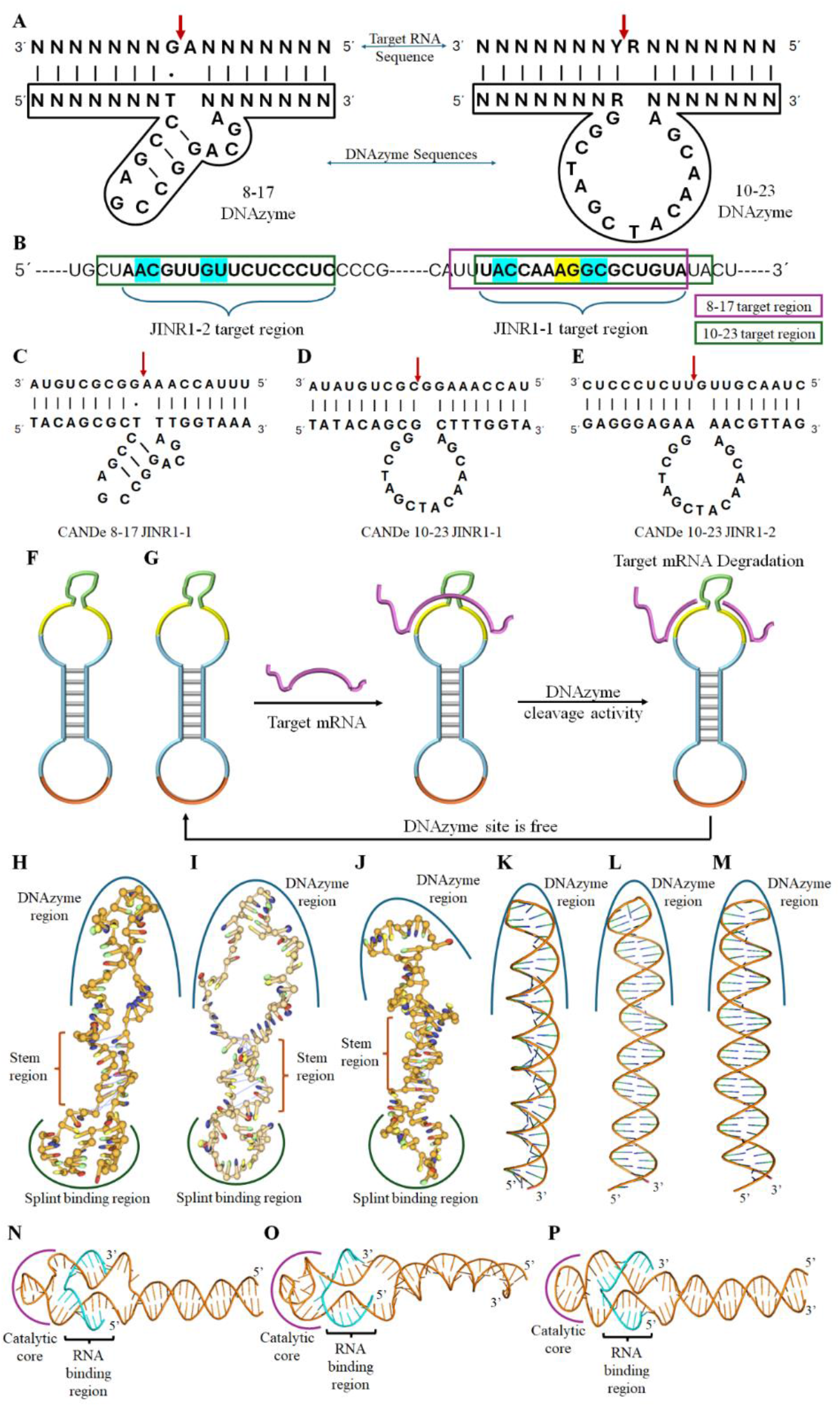
Design, structure, and functioning of CANDe, A, Sequences and secondary structures of 8–17 DNAzyme (left) and 10–23 DNAzyme (right; R = A or G; Y = U or C). B, lncRNA JINR1 sequence with LNA ASO binding sites (JINR1-1, JINR1-2), 8–17 DNAzyme sites (yellow), and 10–23 DNAzyme sites (blue); CANDe target sites for 8–17 (purple box) and 10–23 (green box) indicated. C–E, CANDe sequences showing binding regions and cleavage sites (red arrows): CANDe 8–17 JINR1-1 (C), CANDe 10–23 JINR1-1 (D), CANDe 10–23 JINR1-2 (E). F, Schematic diagram representing the structure of CANDe with splint binding region, target complimentary region, catalytic core sequence, and non-specific regions represented in orange, yellow, green, and blue respectively. G, Schematic diagram illustrating the mechanism of action of CANDe and its recycling within the cell. H-J oxDNA-predicted 3D structures of CANDe 8–17 JINR1-1 (H), CANDe 10–23 JINR1-1 (I), and CANDe 10–23 JINR1-2 (J); DNAzyme, stem, and splint-binding regions highlighted in blue, orange, and green, respectively. K-M, AlphaFold-predicted structures of corresponding linear CANDe constructs; DNAzyme region highlighted in blue: 8–17 JINR1-1 (K), 10–23 JINR1-1 (L), 10–23 JINR1-2 (M). N-O, AlphaFold-based RNA–DNAzyme interaction models: CANDe constructs (orange), RNA target region (blue), catalytic core (purple); 8–17 JINR1-1 (N), 10–23 JINR1-1 (O), 10–23 JINR1-2 (P).

To integrate these catalytic motifs into a modular nanostructure, we designed the CANDe constructs with the following components (Figure 1F): (i) the DNAzyme catalytic core, (ii) 8 nt target-complementary flanking arms on both sides, (iii) a short flexible spacer, (iv) an 8 nt self-complementary stem, (v) an additional spacer, and (vi) terminal 8 nt splint-binding regions at both the 5′ and 3′ ends. The target-complementary arms were positioned over the previously reported ASO JINR1-1 and JINR1-2 binding sites to ensure that these experimentally validated accessible regions would support efficient DNAzyme docking and cleavage and then be reused for other target cleavages (Figure 1G)^41^.

This strategic placement allowed direct benchmarking of DNAzyme-based activity against published LNA ASO-mediated knockdown. The oxView simulation further corroborated that the designed CANDe 8-17 JINR1-1 (Figure H and Figure S1 A), CANDe 10-23 JINR1-1 (Figure 1I and Figure S1 B), and CANDe 10-23 JINR1-2 (Figure 1J and Figure S1 C) constructs adopt the intended structural configuration upon circularization^43^. Spacer segments were incorporated to reduce torsional strain associated with the circular architecture and to maintain proper spatial orientation of functional domains. The metastable 8 nt stem permitted to reversibly transition between hybridized (Figure 1I-K) and unhybridized states (Figure 1F-H). Its low thermodynamic stability promoted splint annealing during circularization as well as allowing target (blue) engagement during catalytic activity, by supporting the hybridisation. This dynamic behaviour was intended to enhance both structural stability in solution and substrate accessibility during cleavage.

### Synthesis and characterization of CANDe

Three distinct CANDe constructs targeting lncRNA JINR1—CANDe 8-17 JINR1-1, CANDe 10-23 JINR1-1, and CANDe 10-23 JINR1-2—were synthesized alongside a Circular Scrambled JINR1-S (Control; Supplementary Table 1). Linear DNA precursors underwent sequential enzymatic processing: 5′-phosphorylation, splint-mediated annealing, T4 DNA ligase-catalyzed circularization, and exonuclease digestion to eliminate residual linear species (Figure 2A). Circularization efficiency was rigorously assessed by electrophoresis of 100 ng aliquots from each processing stage on 15% denaturing urea PAGE, visualized by ethidium bromide staining.

**Figure 2.**
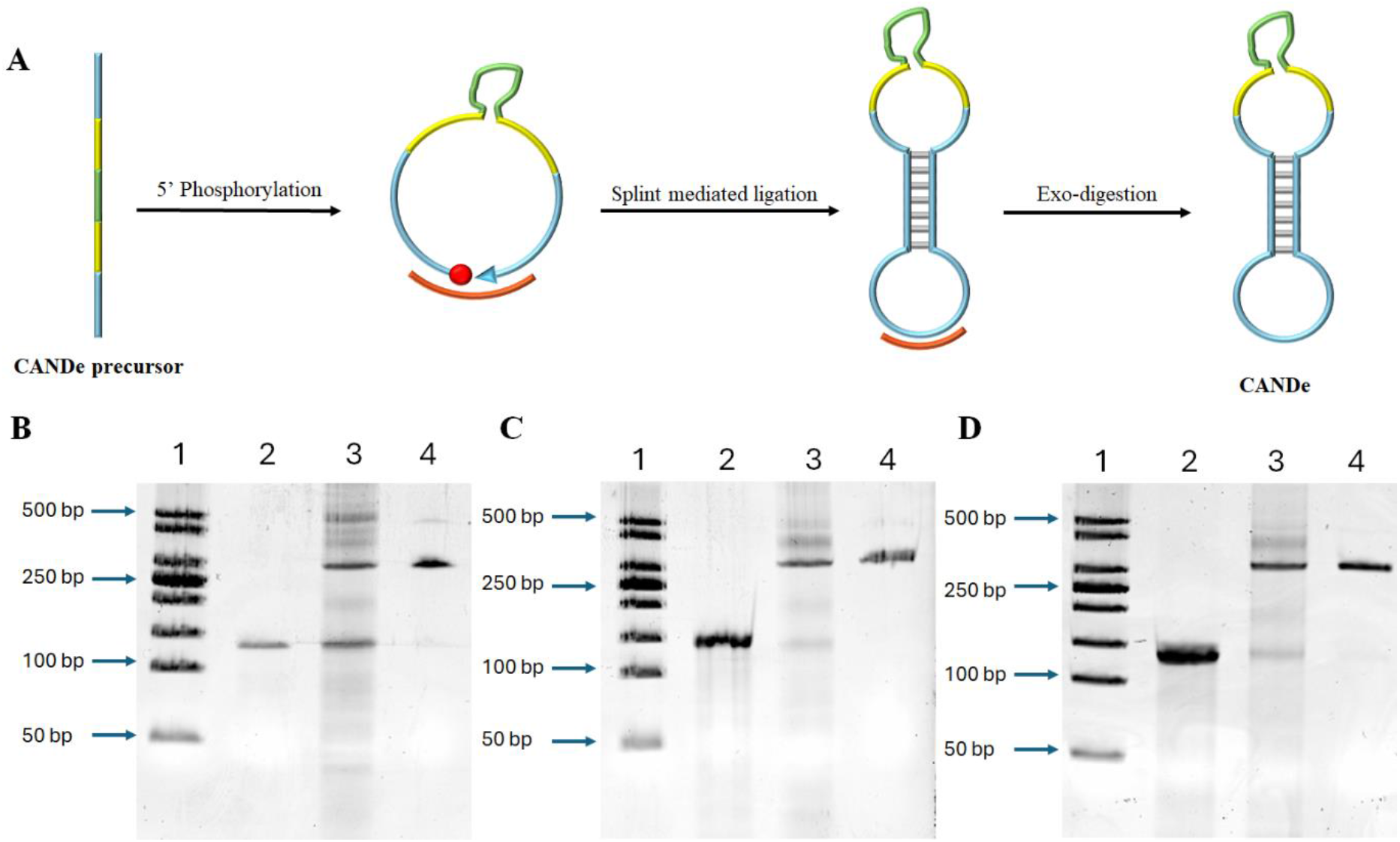
Synthesis of CANDe, A, Schematic diagram represents the steps involved in the synthesis of CANDe highlighting phosphorylation, ligation and exo-digestion steps, B–D, Representative 15% denaturing urea PAGE showing CANDe 8-17 JINR1-1, and CANDe 10-23 JINR1-1, and CANDe 10-23 JINR1-2, respectively, characterisation after different synthesis steps. Lane 1, Ladder (50 bps), Lane 2, untreated control, Lane 3, Ligated sample, Lane 4, Exo digested sample (100 ng each).

For CANDe 8-17 JINR1-1 (Figure 2B), CANDe 10-23 JINR1-1 (Figure 2C), and CANDe 10-23 JINR1-2 (Figure 2D), the linear precursor produced a single, well-defined band in lane 2, serving as the migration reference. Ligation (lane 3) generated a prominent slower-migrating species characteristic of the topologically constrained circular topology, accompanied by faint concatenated species and minor smearing attributable to partial ligation intermediates or adventitious degradation. Critically, the linear precursor band was undetectable post-ligation, confirming near-quantitative end-to-end closure efficiency. Exonuclease treatment (lane 4) then yielded a single, highly discrete resistant band with enhanced electrophoretic mobility relative to the ligated intermediate, unequivocally validating removal of residual linear or nicked species and establishing the structural integrity of the fully circularized CANDe nano-constructs. These data demonstrate robust, reproducible synthesis of nuclease-resistant DNAzyme platforms for therapeutic applications.

### Stability of Linear unmodified DNA v/s CANDe

Oligonucleotide-based therapeutics are highly susceptible to degradation by endogenous nucleases, particularly 3’-exonucleases, due to their rapid hydrolysing nature^44^. They act on the unprotected phosphodiester backbones and compromise therapeutic efficacy in vivo. The Snake Venom Phosphodiesterase (SVPD) assay serves as a robust, standardized model for evaluating nuclease resistance, mimicking physiological degradation pathways to predict the stability of modified oligonucleotides prior to serum or cellular studies^45,46^. To assess the protective efficacy of our nano-construct CANDe, a comparative SVPD stability analysis was performed against unmodified single-stranded DNA (ssDNA). 250 ng samples of each were incubated with SVPD in DMEM media supplemented with 10% FBS at 37°C for 0-90 mins (figure 3A). Degradation profiles were resolved by denaturing urea PAGE, followed by densitometric quantification (Figure 3B).

**Figure 3.**
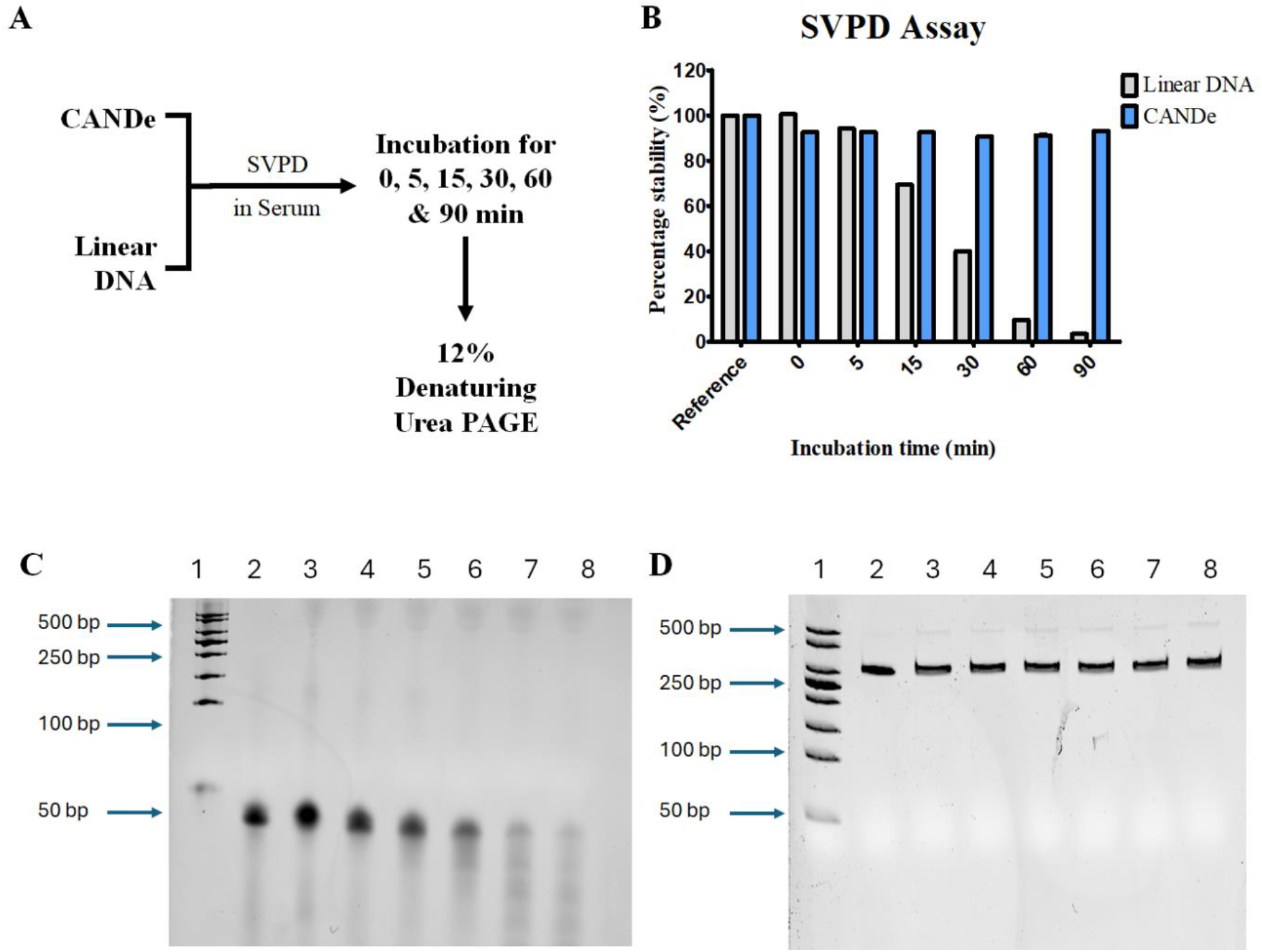
Serum stability assay of CANDe versus linear precursors. **A**, Schematic of the SVPD assay, where samples were incubated with 10% FBS-containing serum for 0–90 min and analysed by 12% denaturing urea PAGE. **B**, Densitometric analysis showing degradation kinetics of linear DNA (grey) and CANDe (blue). **C–D**, Representative 12% denaturing urea PAGE showing stability profiles of linear DNA (panel C), and CANDe (panel D) after SVPD treatment in serum. Lane 1, Ladder (50 bps), Lane 2, untreated control (250 ng), Lane 3-8, 0-, 5-, 15-, 30-, 60-, and 90-min samples respectively (250 ng each).

Unmodified linear DNA underwent rapid degradation upon incubation with snake venom phosphodiesterase in 10% FBS-containing DMEM media. Densitometric analysis revealed a sharp decline in band intensity as early as 5 min (~10% loss; Figure 3C), with only 50% of the intact oligonucleotide remaining after 30 min of treatment (Figure 3B). By 90 min, the band was completely degraded (<5% intact), marked by a progressive increase in smear intensity that confirmed exonucleolytic breakdown across the gel lanes (Figure 3C).

In stark contrast, the CANDe nano-construct remained fully intact across all time points (0–90 min), exhibiting no degradation or smear formation on denaturing urea PAGE (Figure 3D). As a circularized nano-construct lacking backbone modification, CANDe eliminates exposed 3’ or 5’ termini that serve as entry points for exonucleolytic digestion, enabling exceptional evasion of enzymatic hydrolysis. This property preserves >90% structural integrity throughout the assay. These findings underscore CANDe’s topological design advantage for enhancing oligonucleotide therapeutic stability in nuclease-rich physiological environments.

### CANDe Suppresses *JINR1* Expression and Attenuates JEV-Mediated Cell Death

Comparative analysis with LNA ASOs (Figure S2) highlights that CANDe-mediated *JINR1* knockdown yields comparable effects under mock infection conditions. To further evaluate the functional impact of CANDe on JEV-associated lncRNA regulation, SH-SY5Y cells were transfected with CANDe 8-17 JINR1-1, CANDe 10-23 JINR1-1, or CANDe 10-23 JINR1-2, with or without JEV infection, followed by qRT-PCR analysis of JINR1 expression. As shown in Fig. 4B, in mock-infected cells, CANDe 8-17 JINR1-1, CANDe 10-23 JINR1-1, and CANDe 10-23 JINR1-2 reduced *JINR1* expression by ~40%, ~25%, and ~42%, respectively, relative to the JINR1-S (control). JEV infection markedly increased *JINR1* expression (~3-fold) in SH-SY5Y cells transfected with circular scrambled JINR1-S. Importantly, CANDe 8-17 JINR1-1, CANDe 10-23 JINR1-1, and CANDe 10-23 JINR1-2 significantly attenuated this JEV-mediated induction, restricting *JINR1* upregulation to 1.74-, 1.23-, and 1.54-fold, respectively. These results indicate that all three constructs effectively suppressed *JINR1* expression in both mock and JEV-infected cells, with CANDe 10-23 JINR1-1 showing the strongest knockdown efficiency.

**Figure 4.**
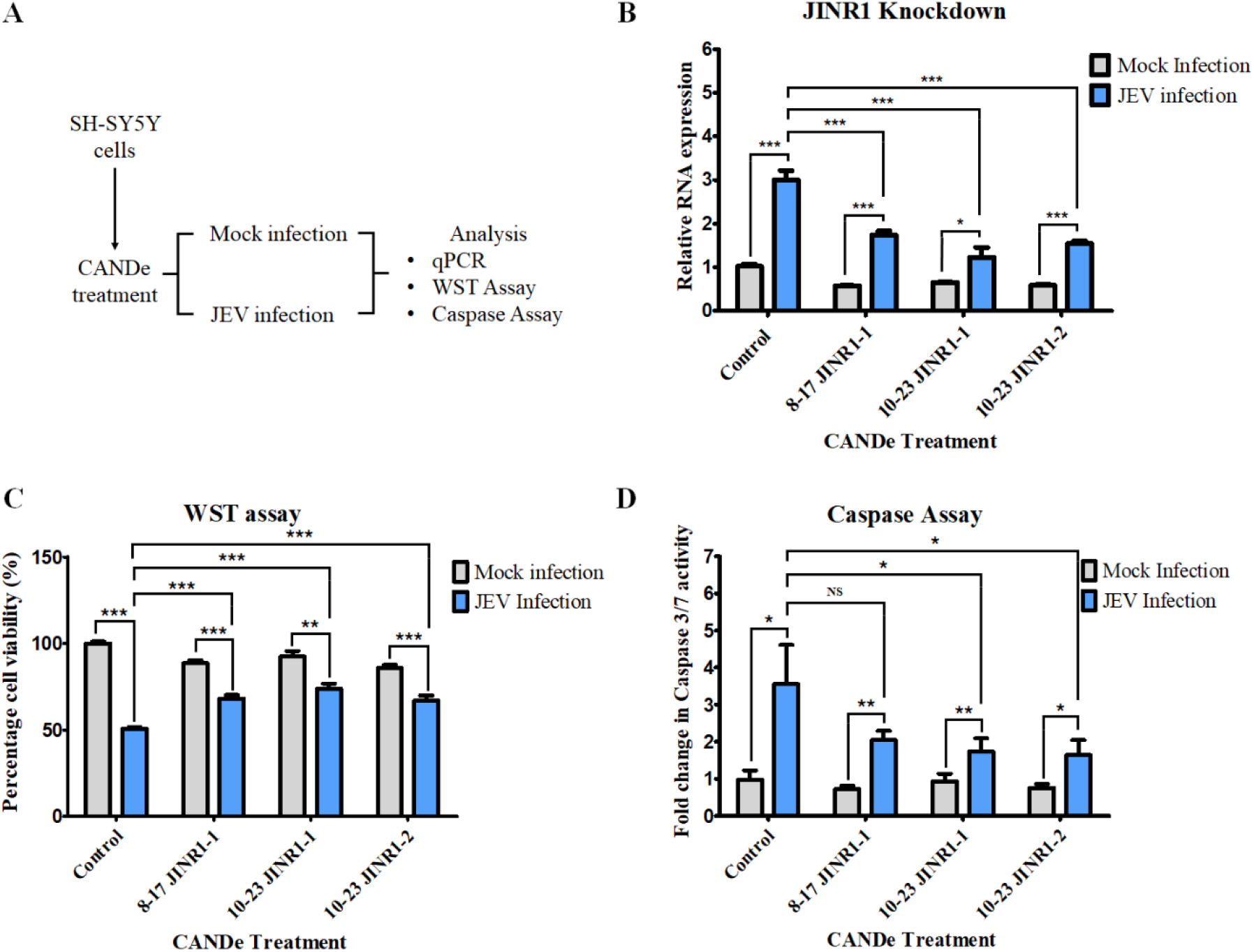
CANDe mediated JINR knockdown and its consecutive effect on JEV-mediated cell proliferation and apoptotic activity. A, Schematic representation of the experimental workflow. SH-SY5Y cells were reverse-transfected with circular scrambled JINR1-S (control), CANDe 8-17 JINR1-1, CANDe 10-23 JINR1-1 or CANDe 10-23 JINR1-2, and subsequently infected with JEV after reaching confluency. Total RNA was isolated 48 h post-infection (hpi), converted into cDNA, and subjected to qRT-PCR analysis; B, qRT-PCR analysis showing relative *JINR1* expression following NaFASO treatment in Mock and JEV-infected cells; C, Proliferation of SH-SY5Y cells measured by WST-1 assay at 48 hpi following *JINR1* knockdown, showing the percentage of viable cells treated with circular scrambled JINR1-S (control), CANDe 8-17 JINR1-1, CANDe 10-23 JINR1-1 or CANDe 10-23 JINR1-2. D, Apoptosis in SH-SY5Y cells measured using the Caspase 3/7 activity at 48 hpi following JINR1 knockdown, showing fold change in Caspase 3/7 activity in cells after treatment with circular scrambled JINR1-S (control), CANDe 8-17 JINR1-1, CANDe 10-23 JINR1-1 or CANDe 10-23 JINR1-2. Hpi hours post-infection. Error bars (n = 3) represent standard deviation. Statistical analysis performed using Student’s t-test. P < 0.05 (*), P < 0.01 (**), P < 0.001 (***), NS, non-significant.

To assess the cytotoxicity and therapeutic potential of CANDe-mediated *JINR1* knockdown, both WST-1 (metabolic activity–based proliferation assay) and Caspase-3/7 (apoptosis assay) analyses were performed. In the absence of infection, CANDe 8-17 JINR1-1, CANDe 10-23 JINR1-1, and CANDe 10-23 JINR1-2 caused only modest reductions in cell proliferation (11.54%, 7.55%, and 14.3%, respectively) in the WST-1 assay, indicating minimal off-target cytotoxicity under basal conditions (Figure 3C). In contrast, JEV infection alone markedly impaired cell proliferation, reducing it to 50.43% of control levels, reflecting the pronounced cytopathic effects of the virus. Notably, CANDe treatment substantially rescued this growth inhibition. CANDe 8-17 JINR1-1 restored proliferation to 88.46%, CANDe 10-23 JINR1-1 to 92.45%, and CANDe 10-23 JINR1-2 to 85.7%. Among these constructs, CANDe 10-23 JINR1-1 demonstrated the strongest protective effect, nearly restoring proliferation to basal levels despite ongoing viral infection.

Consistent with these findings, Caspase-3/7 assay results further supported the protective role of CANDe-mediated *JINR1* knockdown. Under basal conditions, CANDe-treated cells did not exhibit a significant increase in caspase activity relative JINR1-S, indicating that the knockdown constructs did not intrinsically induce apoptotic signaling (Figure 3D). However, JEV infection alone resulted in a marked increase in Caspase-3/7 activity, consistent with enhanced apoptosis. Importantly, CANDe-mediated silencing of *JINR1* significantly attenuated JEV-induced caspase activation, demonstrating a reduction in virus-induced apoptotic cell death. Among the constructs, CANDe 10-23 JINR1-1 again exhibited the most pronounced anti-apoptotic effect, consistent with its superior ability to restore cell proliferation.

Collectively, these findings indicate that CANDe-mediated *JINR1* knockdown mitigates JEV-induced growth suppression and protects neuronal cells from apoptosis. The concordant improvements in cell proliferation and survival strongly suggest that *JINR1* contributes to JEV-induced cytopathology.

### Effect of *JINR1* Knockdown on JEV Replication and GRP78 Expression

To determine whether *JINR1* knockdown influences JEV replication, viral RNA levels and infectious viral titters were evaluated in SH-SY5Y cells transfected with CANDe 8-17 JINR1-1, CANDe 10-23 JINR1-1, or CANDe 10-23 JINR1-2 following JEV infection. Viral RNA levels were quantified using quantitative reverse transcription PCR (qRT-PCR). As shown in Figure 5A, treatment with CANDe 8-17 JINR1-1, CANDe 10-23 JINR1-1, and CANDe 10-23 JINR1-2 resulted in reductions of viral RNA expression by ~35%, ~44%, and ~44%, respectively, compared with the JINR1-S.

**Figure 5.**
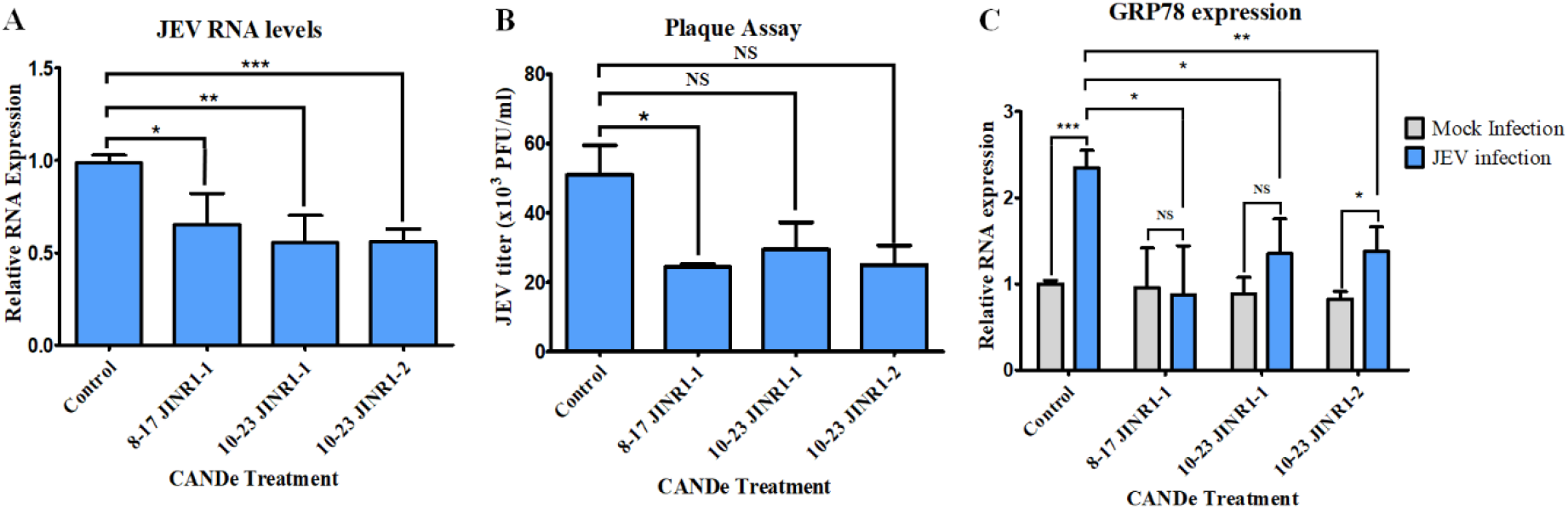
Effect of CANDe-Mediated JINR1 Knockdown on JEV Replication and GRP78 (gene facilitating viral entry). A, Quantitative real-time PCR (qRT-PCR) analysis of Japanese encephalitis virus (JEV) RNA levels following JINR1 silencing. B, Quantification of viral titers expressed as plaque-forming units per milliliter (PFU/mL), demonstrating the effect of JINR1 knockdown on JEV replication efficiency. Relative qRT-PCR expression of the downstream target gene GRP78 in cells treated with CANDe constructs in the presence or absence of JEV infection. Error bars (n = 3) represent standard deviation. Statistical analysis performed using Student’s t-test. P < 0.05 (*), P < 0.01 (**), P < 0.001 (***), NS, non-significant

To further assess the impact of *JINR1* depletion on viral propagation, plaque-forming unit (PFU) assays were performed to quantify infectious virus production. Control cells transformed with JINR1-S and subsequently infected with JEV displayed substantial plaque formation, yielding a mean viral titer of 5.1 × 10^4^ PFU/mL (Figure 5B and Supplementary Figure S3), confirming efficient viral replication under these conditions. In contrast, cells treated with JINR1-targeting CANDe constructs exhibited reduced plaque formation, indicating impaired viral production.

Among the constructs, CANDe 10-23 JINR1-1 reduced viral titers to 2.85 × 10^4^ PFU/mL, corresponding to an approximate 44% decrease relative to JINR1-S. CANDe 8-17 JINR1-1 and CANDe 10-23 JINR1-2 also significantly reduced viral output, yielding titers of 2.45 × 10^4^ PFU/mL and 2.5 × 10^4^ PFU/mL, respectively, representing ~51% reductions relative to JINR1-S. These findings suggest that *JINR1* functions as a proviral host factor whose silencing impairs JEV replication and production of infectious virions. Overall, the CANDe constructs exhibited comparable antiviral activity, with modest variations likely reflecting differences in target accessibility or cleavage efficiency within the *JINR1* transcript.

GRP78 is known to facilitate viral entry, promote replication, and support endoplasmic reticulum stress adaptation during JEV infection^41,42,47^. Previous studies by Tripathi et al. demonstrated that lncRNA-*JINR1* enhances JEV replication by regulating GRP78 expression^42,^. Therefore, we examined the effect of CANDe-mediated *JINR1* knockdown on GRP78 expression following JEV infection. As shown in Figure 5C, in mock-infected cells, CANDe 8-17 JINR1-1, CANDe 10-23 JINR1-1, and CANDe 10-23 JINR1-2 reduced GRP78 expression by ~5%, ~40%, and ~20%, respectively, relative to JINR1-S.

Upon JEV infection, cells transfected with the JINR1-S exhibited a 2.35-fold induction of GRP78 expression. In contrast, CANDe 8-17 JINR1-1, CANDe 10-23 JINR1-1, and CANDe 10-23 JINR1-2 limited this induction to 0.87-, 1.35-, and 1.38-fold, respectively. These results indicate that *JINR1* knockdown suppresses GRP78 upregulation during JEV infection. Taken together, these findings establish a functional link between CANDe-mediated *JINR1* silencing, reduced GRP78 expression, and inhibition of JEV replication, highlighting the potential of targeting *JINR1* as a strategy for limiting JEV-induced cellular pathology.

## Discussion

Japanese encephalitis virus (JEV), a neurotropic member of the genus *Orthoflavivirus* (formerly *Flavivirus*), remains a major public health concern across South and Southeast Asia and has recently expanded into northern regions of Australia^48^. Despite the availability of licensed vaccines—including inactivated Vero cell–derived and live attenuated formulations— no specific antiviral therapy is currently approved for clinical management of JEV infection^32–35^. Vaccination programs have significantly reduced disease incidence; however, incomplete vaccine coverage, waning immunity, and viral ecological persistence continue to permit outbreaks. Clinical management therefore remains largely supportive, underscoring the need for therapeutic strategies capable of mitigating viral replication and virus-induced neuropathology.

Host-directed antiviral approaches have gained attention as a complementary strategy to traditional pathogen-targeted interventions^49^. In this context, the long noncoding RNA (lncRNA) *JINR1* has emerged as a proviral host factor implicated in JEV infection. Antisense oligonucleotide (ASO)-mediated silencing of *JINR1* has demonstrated effective suppression of viral propagation, highlighting this transcript as a promising therapeutic target^41,42^. However, conventional ASOs function primarily through recruitment of endogenous RNase H1 or through steric blockade mechanisms, thereby relying on host cellular machinery for RNA degradation or translational inhibition^9^. Such dependence may influence efficiency across cell types and physiological conditions, particularly in virally stressed environments where host pathways are dysregulated.^50,51^ Moreover, ASOs typically require extensive backbone modifications to enhance nuclease resistance, which can increase manufacturing complexity and potentially affect pharmacodynamic properties.

Catalytic DNAzymes offer a mechanistically distinct alternative. Unlike ASOs, DNAzymes mediate direct, sequence-specific cleavage of RNA substrates through intrinsic catalytic activity and do not require host enzymatic machinery for target degradation^15^. This self-sufficient mode of action enables catalytic turnover, theoretically allowing multiple RNA cleavage events per DNAzyme molecule and potentially reducing the intracellular concentration required for effective knockdown. Nevertheless, broader therapeutic translation of DNAzymes has historically been limited by rapid exonucleolytic degradation and the need for stabilizing chemical modifications that may compromise catalytic efficiency, particularly through perturbation of metal-ion coordination within the catalytic core.

In the present study, we addressed these limitations through the development of Circular Active Nano DNAzymes (CANDe), an enzymatically synthesized circular DNAzyme platform designed to achieve structural stabilization without backbone modification. Circularization eliminates exposed 3′ and 5′ termini, the primary entry points for exonucleases, thereby enhancing resistance to degradation while preserving catalytic integrity. The CANDe constructs were rationally engineered to incorporate discrete functional modules: splint-binding domains enabling efficient ligation, a stem region providing structural organization, substrate-recognition arms mediating target hybridization, and a catalytically active DNAzyme core. This modular architecture permits maintenance of catalytic accessibility while achieving topological stabilization. Importantly, stabilization is achieved through architecture rather than chemical alteration, preserving native catalytic functionality.

Topological circularization conferred pronounced resistance to nuclease-mediated degradation under exonuclease-rich conditions in which linear DNAzymes were rapidly degraded. These findings demonstrate that architectural engineering can substitute for backbone chemical modification in enhancing DNAzyme stability. From a translational perspective, this approach simplifies synthesis, reduces reliance on modified phosphoramidites, and may facilitate scalable production. These results suggest that architectural engineering can substitute for chemical modification in enhancing DNAzyme stability, with important implications for scalability, cost, and regulatory translation.

Functionally, CANDe-mediated silencing of *JINR1* resulted in robust attenuation of JEV-induced cytopathology in neuronal cell models. Treatment restored cellular viability and significantly reduced caspase-3/7 activation in infected cells, indicating mitigation of virus-induced apoptotic pathways. Importantly, no detectable cytotoxicity was observed in uninfected cells, supporting the specificity and therapeutic window of host-factor targeting. Given that neuronal loss underlies the severe and often irreversible neurological sequelae associated with Japanese encephalitis, preservation of cell viability is a particularly meaningful outcome.

At the molecular level, CANDe treatment produced significant downregulation of *JINR1* expression, confirming efficient intracellular catalytic cleavage. This suppression was accompanied by reduced expression of GRP78, a host chaperone implicated in orthoflavivirus replication and endoplasmic reticulum stress adaptation. Concomitant decreases in viral RNA copy number and infectious viral titers further established a mechanistic link between DNAzyme-mediated host RNA cleavage and impaired viral propagation. Plaque assays independently validated these findings, demonstrating marked reductions in viral yield following CANDe treatment.

Comparative analysis of distinct CANDe variants revealed that both DNAzyme motif selection and target-site positioning critically influence antiviral efficacy. Constructs incorporating either the 8–17 or 10–23 catalytic cores exhibited functional activity; however, the 10–23-based CANDe targeting the JINR1-1 site consistently produced superior transcript knockdown, viral suppression, and cell survival outcomes. These observations underscore the importance of integrating catalytic motif properties with RNA structural accessibility during rational DNAzyme design.

The host-directed strategy described herein offers advantages over direct viral targeting approaches. RNA viruses such as JEV exhibit high mutation rates, facilitating escape from therapeutics directed against viral proteins or genomic RNA. In contrast, targeting a relatively conserved host factor reduces the probability of resistance arising through viral genetic variation. Additionally, the catalytic turnover intrinsic to DNAzymes may enable effective silencing at lower doses compared to stoichiometric antisense platforms.

Beyond the immediate antiviral application, the CANDe platform possesses inherent modularity. The substrate-recognition arms can be readily redesigned to target alternative RNA sequences while retaining the same catalytic core and circular architecture. This adaptability enables rapid retargeting toward other viral RNAs, host dependency factors, oncogenic transcripts, or inflammatory mediators. Because stabilization is conferred by circular topology rather than sequence-specific chemical modification, redesign does not necessitate re-optimization of backbone chemistry, thereby streamlining development pipelines.

From a manufacturing standpoint, the enzymatic splint-mediated circularization strategy represents a scalable and potentially more sustainable alternative to fully synthetic modified oligonucleotides. High circularization efficiency and structural homogeneity support feasibility for further translational development. Nevertheless, comprehensive evaluation of in vivo pharmacokinetics, biodistribution, immunogenicity, and delivery strategies will be essential to determine clinical potential.

## Conclusion

This study introduces Circular Active Nano DNAzymes (CANDe) as a structurally engineered and catalytically active platform for host-directed antiviral intervention against Japanese encephalitis virus (JEV). By targeting the proviral lncRNA *JINR1*, CANDe achieves efficient RNA cleavage, resulting in suppression of viral replication and attenuation of virus-induced cytopathology in neuronal cells. In contrast to antisense oligonucleotides, which rely on endogenous cellular machinery for target degradation, CANDe mediates direct and sequence-specific RNA cleavage through intrinsic catalytic activity. This mechanistic independence is particularly advantageous in virally stressed environments where host pathways may be dysregulated. Topological circularization enhances nuclease resistance by eliminating exposed termini, thereby improving structural stability without requiring backbone chemical modifications that can compromise catalytic efficiency or increase synthetic complexity. The combination of preserved catalytic function and enhanced stability highlights architectural engineering as an effective alternative to conventional chemical stabilization strategies.

The modular design of CANDe further enables rapid retargeting through simple redesign of substrate-recognition arms, facilitating application against diverse viral RNAs, host dependency factors, or disease-associated transcripts. Future work should prioritize in vivo evaluation of pharmacokinetics, biodistribution, and delivery strategies, particularly for neurotropic infections. Structural optimization, including modulation of stem architecture and hybridization parameters, may further refine the balance between stability and catalytic turnover. Overall, CANDe establishes circular DNAzymes as a scalable and versatile modality for catalytic gene silencing, expanding the translational landscape of nucleic acid therapeutics for antiviral and broader RNA-targeted applications.

## Author contributions

CS, SS, VS, and SG designed the study. CS, SS, and DS performed the experiments. CS, SS, and DS performed the data analysis. CS, SS, VS, and SG wrote the manuscript. All authors edited and approved the final manuscript.

## Data Availability

The data for this manuscript is available upon reasonable request.

## Acknowledgement

This work was supported by the DST-Anusandhan National Research Foundation (ANRF)-CRG/2023/000822 to VS, and Mahindra University internal funding. The authors are grateful to Prof. Rajinder S. Chauhan (Dean, Centre for Life Sciences) and Prof Yajulu Medury (Vice Chancellor, Mahindra University) for their constant scholarly encouragement. We acknowledge the use of the grammar correction tool “Grammarly” in editing the manuscript. SS is supported by CSIR SRF fellowship (File no. 09/1026(23714/2025-EMR-I)

## Electronic Supplementary Information (ESI)

Oligonucleotide sequences, CANDe secondary structures, and plaque assay representative images.

