## Supplemental Data for "Targeting lncRNA *JINR1* with programmable Circular Active Nano DNAzyme (CANDe) suppresses Japanese Encephalitis Virus infection"

**Supplementary Table 1:** List of sequences of various oligonucleotides used in different experiments

| Name | Sequence (5'-3') (italics: splint binding region; underlined: target binding region; bold: DNAzyme ) |
|---|---|
| CANDe 8-17 JINR1-1 | *GACCTGCTA*TAGTAGGGACTGAAAATACAGCGC**TCCGAGCCGGACGA**TTGGTAAAAAAACAGTCCCATGAT*CTGGACGAG* |
| CANDe 10-23 JINR1-1 | *GACCTGCTA*TAGTAGGGACTGAAAATATACAGC**GGGCTAGCTACAACGA**CTTTGGTAAAAACAGTCCCATGAT*CTGGACGAG* |
| CANDe 10-23 JINR1-2 | *GACCTGCTA*TAGTAGGGACTGAAAAGAGGGAG**AGGCTAGCTACAACGA**AACGTTAGAAAACAGTCCCATGAT*CTGGACGAG* |
| Circular scrambled JINR1-S (Control) | *GACCTGCTA*GACTGAAAACGGAAAGAGTGCGATAAAACAGTC*CTGGACGAG* |
| Splint | TAGCAGGTCCTCGTCCAG |
| LNA JINR1-1 | TACAGCGCCTTTGGTA |
| LNA JINR1-2 | GAGGGAGAACAACGTT |
| Scrambled LNA | AACACGTCTATACGC |
| JINR1 fwd primer | CAGTGACGGAACAGTACCAG |
| JINR1 rev primer | TCACAAACATCCCGCTCT |
| GRP78 fwd primer | CTGTCCAGGCTGGTGTGCTCT |
| GRP78 rev primer | CTTGGTAGGCACCACTGTGTTC |
| JEV fwd primer | GAGCTTGTTGGACGGCAGAG |
| JEV rev primer | CACGGCGTCGATGAGTGTTC |

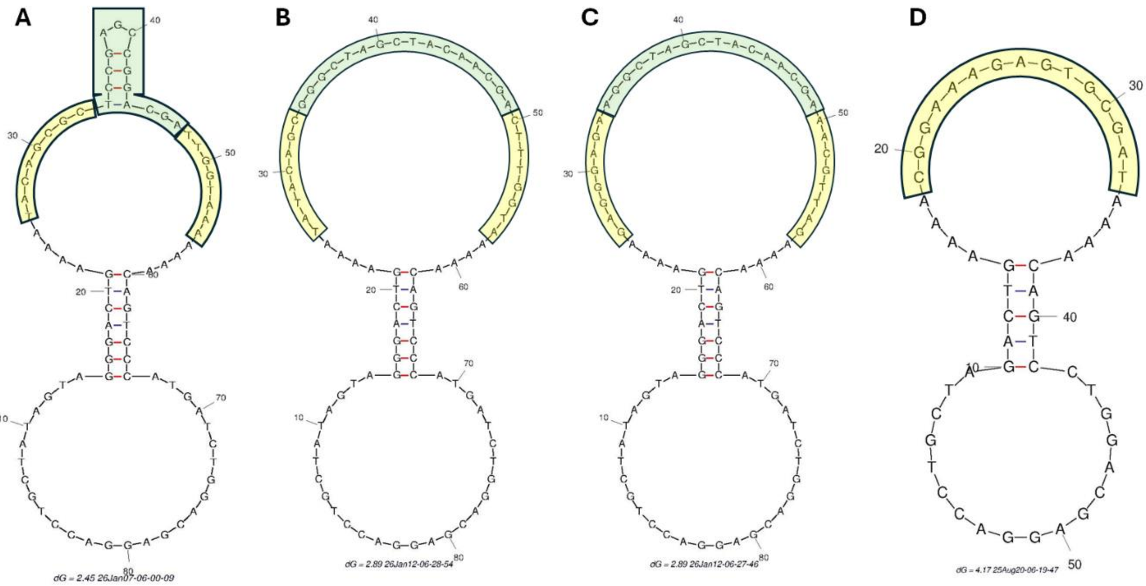


**Figure S1.** Secondary structure of CANDe. A, mFold image of CANDe 8-17 JINR1-1 sequence with DNAzyme catalytic core sequence and binding arms highlighted in green and yellow respectively. B, mFold image of CANDe 10-23 JINR1-1 sequence with DNAzyme catalytic core sequence and binding arms highlighted in green and yellow respectively. C, mFold image of CANDe 10-23 JINR1-2 sequence with DNAzyme catalytic core sequence and binding arms highlighted in green and yellow respectively. D, mFold image of Circular Scrambled JINR1-S with the scrambled binding arm sequence highlighted in yellow.

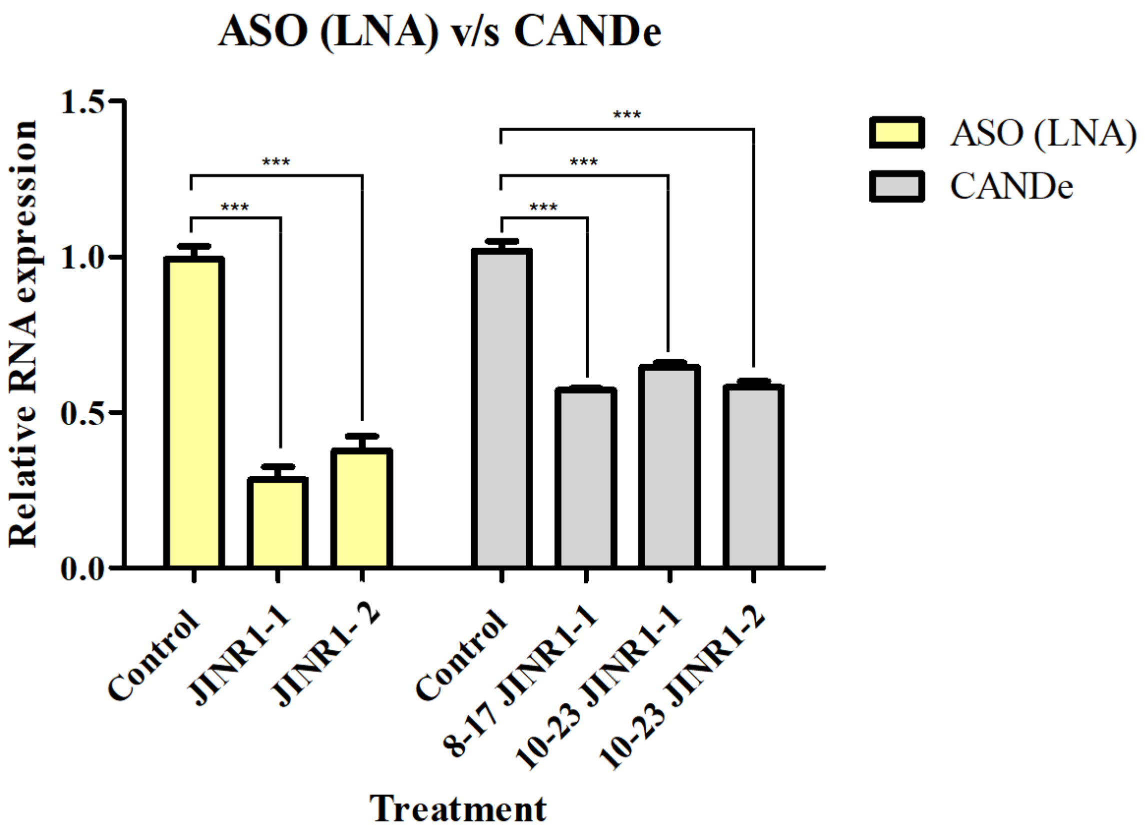


**Figure S2.** Comparative Efficiency of LNA-ASO and CANDe–Mediated *JINR1* Knockdown in SH-SY5Y Cells. Bar graph showing comparative analysis of CANDe- and LNA-ASO–mediated *JINR1* knockdown under basal (uninfected) conditions in SH-SY5Y cells, assessing the relative silencing efficiency of the two antisense strategies. Error bars (n = 3) represent standard deviation (SD). Statistical significance determined using Student's t-test. $P < 0.05$ (*), $P < 0.01$ (**), $P < 0.001$ (***); NS, not significant.

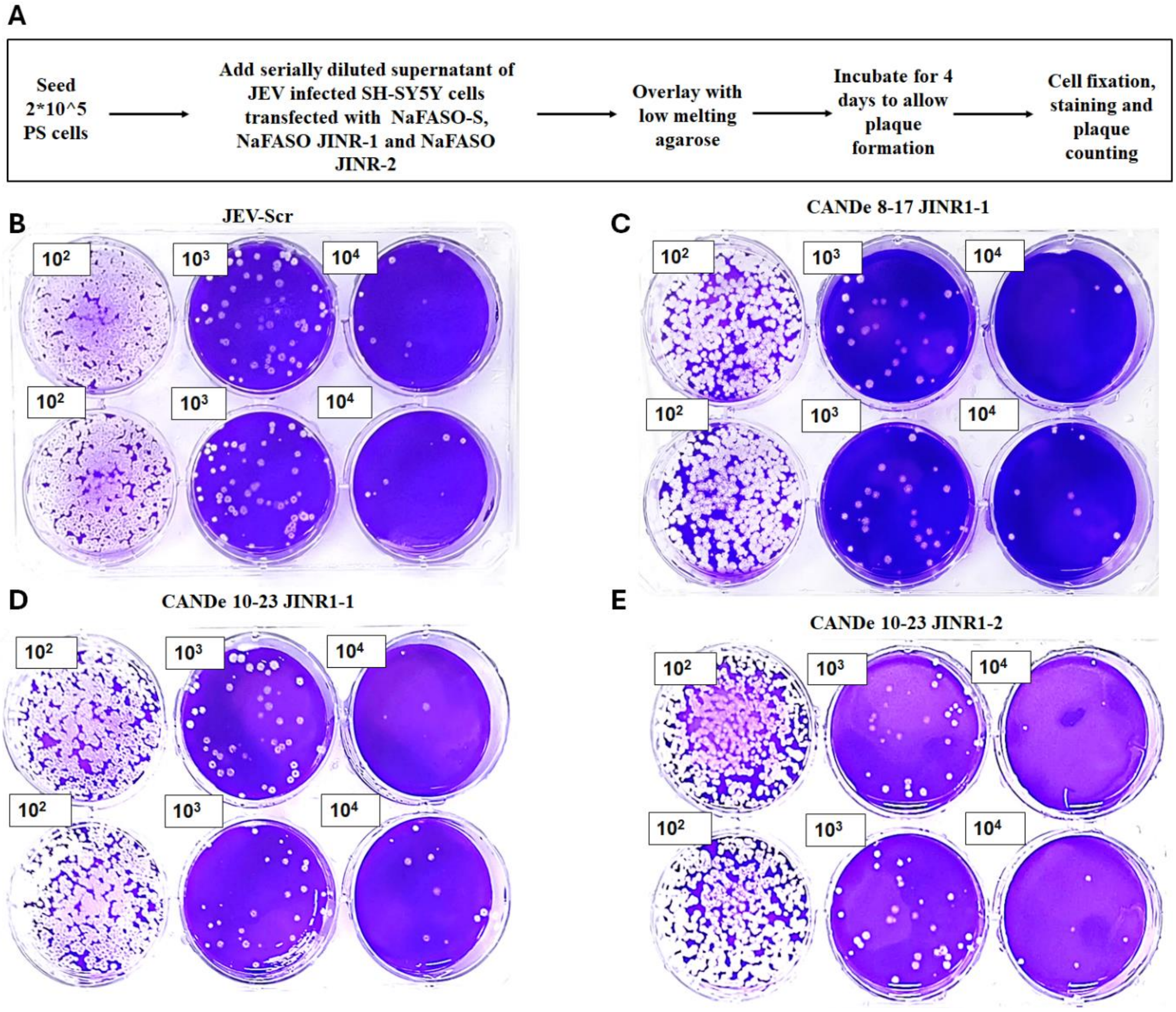


**Figure S3.** Plaque assay workflow and representative results from different samples. A. Schematic representation of the experimental workflow used for the JEV plaque assay. B-E, Representative images of plaque assays performed using JEV viral samples from cells treated with circular scrambled JINR1-S (control), CANDe 8-17 JINR1-1, CANDe 10-23 JINR1-1, and CANDe 10-23 JINR1-2 respectively, showing variations in density corresponding to viral infectivity at different viral titter dilutions.
